# Routine FFPE sections support clinically compatible single-nucleus transcriptomics across six human cancer types

**DOI:** 10.64898/2026.07.23.740343

**Authors:** Jasper Wouters, Juliette Bertorello, Marine Gaillard, Baptiste Simon, Sarah Gastineau, Amélie Roehrig, Maud Dupont-Roc, Elise Amblard, Amanda Pupo, Helena Yu, Jean-Yves Blay, Coralie Guérin, Laetitia Nebot-Bral, Anne Vincent Salomon, Loïc Verlingue, Evanguelos Xylinas, Luc Cabel, Jeffrey S. Ross, Vincent Miller, Eric Letouze, Celine Vallot

**Author notes:** co-first authors (Jasper Wouters, Juliette Bertorello, Marine Gaillard). co-last authors (Eric Letouze, Celine Vallot).

## Abstract

Tumor cellular composition—including malignant cell states, immune populations, and stromal populations—is increasingly recognized as a determinant of therapeutic response and resistance to anti-cancer agents, yet comprehensive cellular profiling remains largely confined to research settings. Here, we present a clinically compatible sample-to-report workflow for tumor composition profiling from routine formalin-fixed paraffin-embedded (FFPE) clinical specimens. By combining low-input single-nucleus RNA sequencing with foundation model-based automated cell annotation, this workflow enables prospective sample-by-sample analysis without dedicated research material or cohort-based processing. Across 116 clinical specimens representing six cancer types, we generated reproducible measurements of cellular composition and cell-type-specific gene expression, demonstrated high technical reproducibility, and showed concordance with pathological assessment of immune infiltration. The workflow was similarly applicable to archival FFPE material and ultra-low-input biopsy specimens. Together, these findings establish a practical framework for routine single-cell profiling from standard pathology specimens and open the perspective of prospective evaluation of cellular composition as a clinical biomarker in precision oncology.

## Introduction

Intratumoral cellular and genomic heterogeneity is increasingly recognized as a major determinant of clinical outcome in cancer^1–3^. Distinct malignant, immune, and stromal cell populations can coexist within the same tumor and influence therapeutic response and the emergence of treatment resistance. Current clinical assays partially capture this complexity only by bulk measurement where the average signals across all cells present in a specimen is reported and therefore only a partial view of tumor biology is provided. In addition, pathology-based approaches provide cellular resolution but are typically limited to a predefined set of expressed proteins and other markers. Cellular profiling approaches based on single-cell transcriptomics address these limitations by enabling transcriptome-wide characterization of individual malignant, immune, and stromal populations within the tumor ecosystem^4^. Across multiple cancer types, single-cell studies have identified in patients clinically relevant tumor states^5–7^, mechanisms of resistance^8,9^, and microenvironmental determinants of therapeutic response^10–13^ that are not captured by conventional approaches.

Despite previously demonstrated utility in investigative settings, there has been only very limited penetrance of single-cell based assays in clinical workflows^14,15^. Most clinical studies continue to utilize fresh or fresh-frozen tissues^16,17^, requiring dedicated and rapid tissue processing, as well as substantial amounts of starting material. These requirements contrast with typical clinical formalin-fixed paraffin embedded (FFPE) tissue block samples, which often are limited in overall volume and in the proportion of tumor versus benign cells. In addition, most studies are structured as cohorts, with the summation of multiple patient samples, limiting the ability to function on a sample-per-sample basis, a necessity for prospective patient profiling.

To address limitations in sample conservation, several FFPE-compatible transcriptomic technologies have emerged. Spatial transcriptomic approaches^18,19^ preserve tissue architecture and provide complementary information on tissue organization and cell-cell interactions, whereas single-nucleus RNA sequencing enables comprehensive transcriptome-wide characterization of individual cellular populations. Alternatively, recently developed FFPE single-nucleus RNA-sequencing methods enable transcriptome-wide cellular profiling but typically require dedicated thick tissue curls (requiring more than one 25µm curls) that are often unavailable in routine clinical practice^20–22^. Consequently, a clinically scalable workflow compatible with routine pathology sections remains lacking.

Here, we developed a clinically compatible FFPE single-nucleus RNA-sequencing workflow that operates on four routine 5-μm FFPE pathology sections. By combining low-input FFPE nuclei preparation with foundation model-based cell annotation, the workflow enables scalable sample-by-sample profiling of clinical specimens without requiring cohort-level integration. We validated the workflow for 116 samples, including 36 biopsies, across six major cancer types— bladder, breast, colon, lung, ovarian, and pancreatic carcinomas—demonstrating robust performance, reproducibility, and biological fidelity across these diverse histologies.

## Results

### Development of a clinically applicable FFPE single-nucleus RNA-sequencing workflow across six cancer types

To enable integration of single-cell transcriptomics into routine pathology practice, we developed a complete sample-to-report workflow designed to accommodate routine clinical oncology practice. Unlike research-oriented single-cell studies that typically rely on fresh or frozen tissue, cohort-based processing, and expert-intensive data interpretation, our approach was designed to operate on FFPE specimens individually available during routine patient care using automated, standardized foundation-model–based analysis (**Figure 1A**).

**Figure 1:**
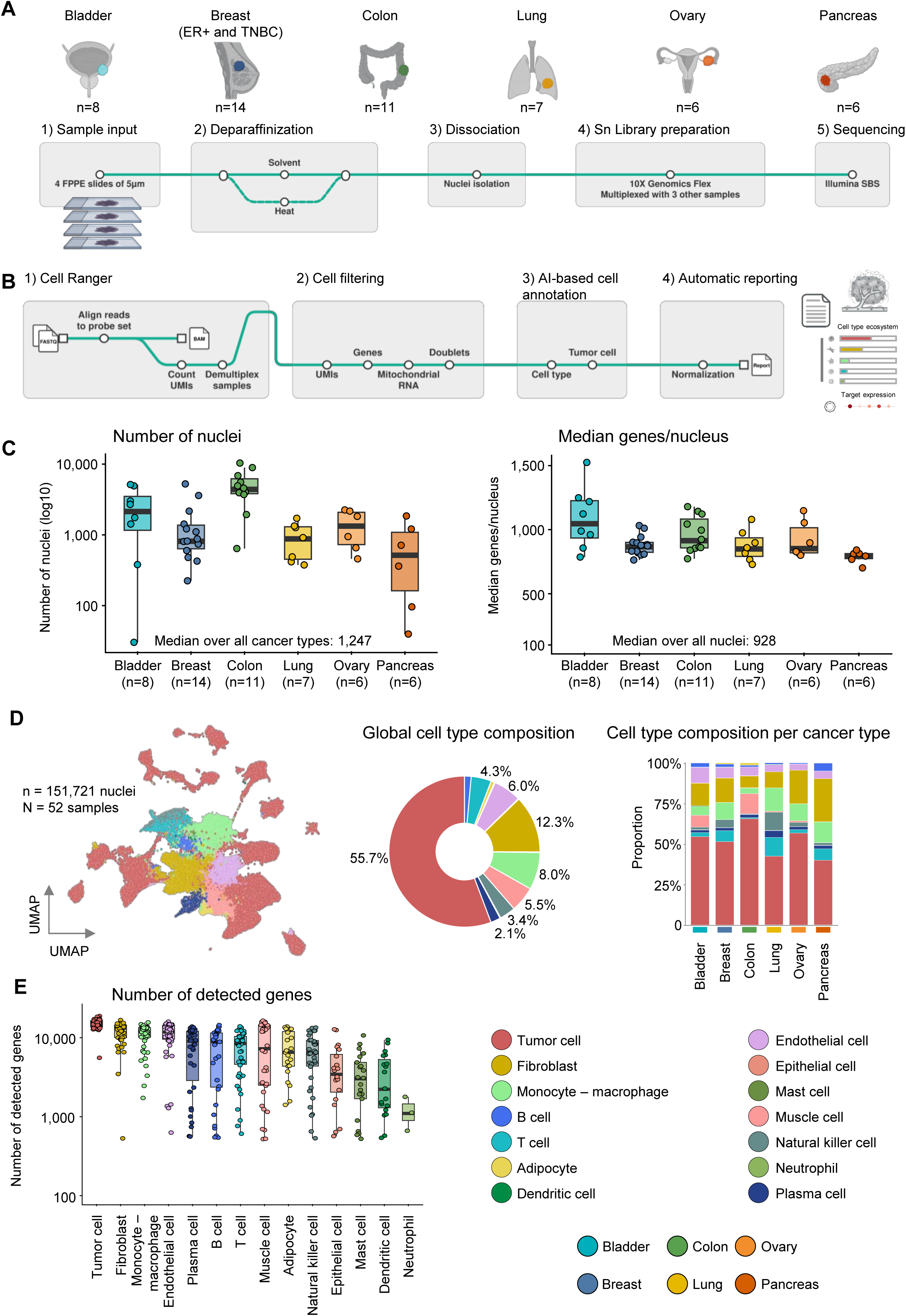
Development of a clinically compatible FFPE cellular profiling workflow. (A) Overview of our multicenter cohort of 52 unique surgical resection specimens. Experimental single-nucleus workflow from FFPE sections. (B) Sample-by-sample workflow, starting from four routine 5-μm pathology sections to an automated report, and including automated cell annotation and target expression quantification. (C) Boxplots representing numbers of nuclei (left) and numbers of genes per nuclei (right) obtained for each resection sample (n=52), grouped by cancer types. (D) Left: UMAP embedding of the single-nucleus dataset of 52 resections, nuclei are colored according to cell type. Middle: general cell type composition of the full dataset. Right: Barplot representing cell type composition by cancer type. (E) Distribution of detected genes per cell type across individual specimens.

The workflow combines a low-input FFPE single-nucleus RNA-sequencing protocol based on the Chromium Single Cell Gene Expression Flex v2 chemistry (10x Genomics) with OneMap, an automated foundation model-based analysis framework for cell annotation and reporting. To preserve diagnostic material, the assay was optimized to utilize four FFPE 5-μm pathology sections. Following sequencing, the OneMap algorithm automatically identifies malignant, immune and stromal populations, quantifies cell-type-specific gene expression, and generates standardized descriptions of tumor composition without requiring cohort-level integration or manual expert annotation (**Figure 1B**).

To evaluate the performance of this workflow in the clinical setting, we assembled a multicenter cohort comprising 88 routine FFPE clinical specimens from 83 patients, collected at Institut Curie, Centre Léon Bérard, Hôpital Bichat–Claude Bernard, and Memorial Sloan Kettering Cancer Center (MSKCC). The cohort included 52 surgical resections and 36 biopsies, spanning six major cancer types: bladder, breast, colon, lung, ovarian, and pancreatic cancer. The breast tumors included both estrogen receptor-positive (ER+) and triple-negative breast cancer (TNBC) cases. In total, 116 single-nucleus RNA-sequencing experiments were performed, including technical replicates used for analytical validation (**Table S1**).

We first characterized assay performance using the 52 surgical resection specimens, which generated a total of 151,721 nuclei (**Figure 1C-D**). We isolated a median number of 1,247 nuclei per specimen, with a median gene coverage of 928 detected genes per nucleus. To enable prospective sample-by-sample analysis, data were processed using OneMap, that leverages scGPT^23^ foundation-model embeddings to classify cellular populations from individual samples. OneMap combines reference-based annotation of normal cell types with a dedicated framework for malignant cell classification and generates standardized reports of tumor composition together with cell-type-resolved expression of clinically relevant therapeutic targets. Unlike conventional single-cell analysis workflows, which typically require aggregated cohort-level integration and manual annotation, OneMap provides a standardized real-time interpretation of individual clinical specimens, making the analytical workflow compatible with prospective patient-level analyses, particularly in the context of clinical trials

Across the cohort, OneMap identified a median of 10 cellular populations per specimen (range, 4–14; **Figure S1A**), demonstrating consistent identification of tumor ecosystem composition across all six cancer types. Overall, 18,523 genes were detected across the cohort. Individual specimens contained a median of 15,524 detected genes. Several thousand genes were detected within each cell type, with the exact number of identified genes depending on both the transcriptional complexity and the abundance of the cell type (**Figure 1E, Figure S1B**). For subsequent analyses, we set the threshold of a successful tumor composition profile to at least 500 nuclei with more than 500 detected genes per nucleus, enabling access to rare cells from the TME e.g NK cells.

Together, these results demonstrate that routine FFPE pathology sections can support standardized cellular profiling across diverse solid tumors using a foundation model–based analysis integrated into a sample-to-report workflow.

### Defining specimen requirements for routine FFPE tumor ecosystem profiling

To establish practical specimen requirements for routine pathology laboratories, we evaluated the impact of key pre-analytical variables on assay performance. To assess the impact of deparaffinization, samples processed with and without solvent treatment (xylene) were compared (n=12; **Figure 2A**). Xylene-treated samples, that have undergone extensive deparaffinization yielded substantially higher numbers of recovered nuclei (median 1,831 versus 619 nuclei, p-value = 3.9×10^-4^), while gene coverage remained comparable between the two approaches (median 801 versus 876 genes per nucleus). Although both workflows generated data with sufficient coverage to decode the tumor micro-environment (TME) and quantify gene expression levels, xylene treatment consistently improved nuclei recovery and was therefore selected for subsequent analyses.

**Figure 2:**
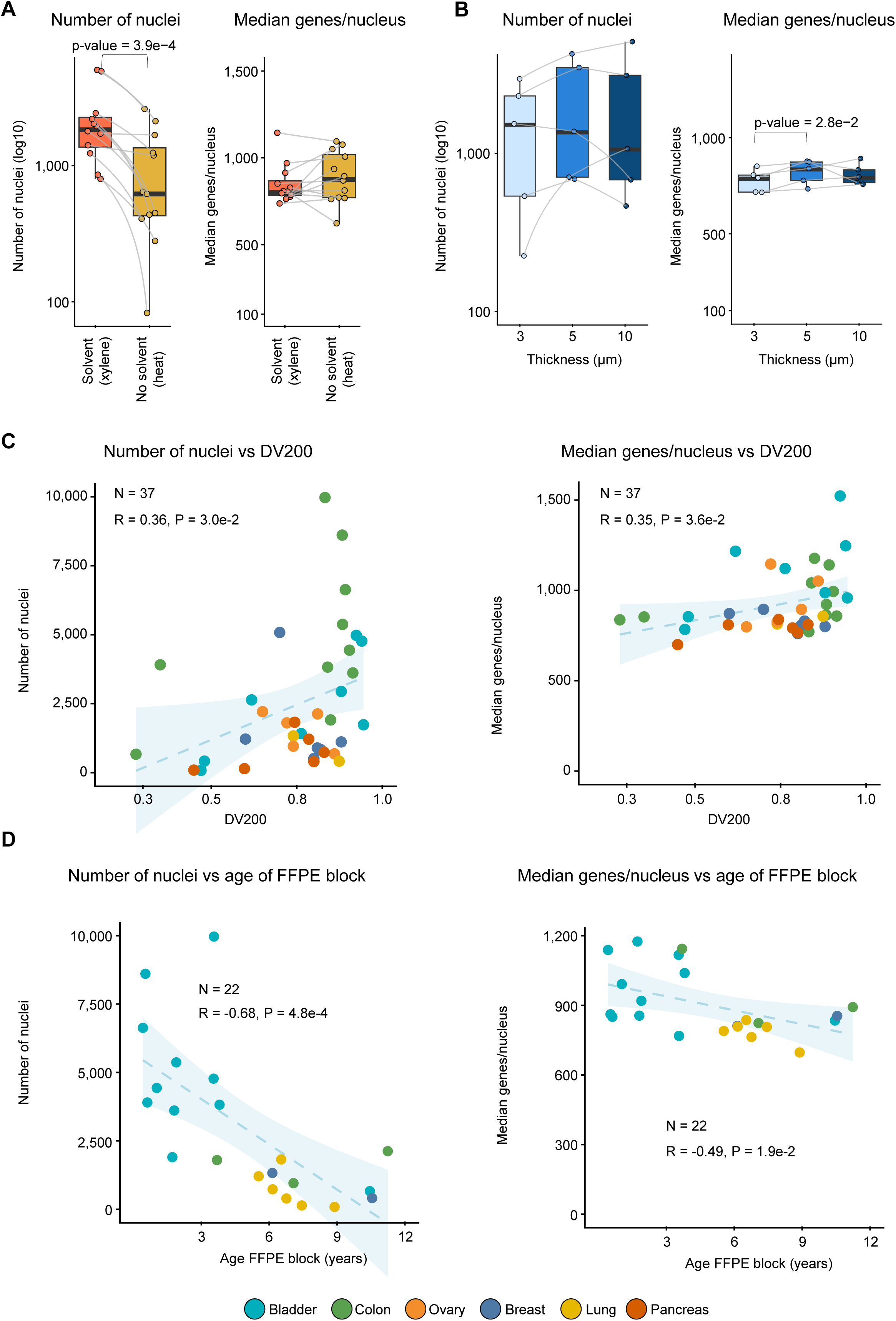
Defining specimen requirements for routine FFPE cellular profiling. (A) Boxplot representations of number of nuclei and median coverage, genes per nucleus, per sample with or without xylene exposure. P-values for paired t-test are displayed when below 0.05. (B) Plots as in A for sections of 3, 5 or 10µm thickness. P-values for paired t-test are displayed when below 0.05. (C) Scatterplot representing number of nuclei and coverage per nuclei versus DV200 values, assessing RNA quality of each sample. Each dot corresponds to one specimen and is colored according to cancer type. The number of samples included in analysis is indicated at the top left corner of graphs, together with Pearson’s R and corresponding p-values. (D) Scatterplot representing number of nuclei and coverage per nuclei versus age of the sample. Each dot corresponds to one specimen and is colored according to cancer type. The number of samples included in analysis is indicated at the top left corner of graphs, together with Pearson’s R and corresponding p-values.

We next established the tissue input requirements compatible with routine pathology practice by evaluating FFPE section thickness (**Figure 2B**). For breast cancer samples (n=5), FFPE sections of 3, 5 and 10 μm were evaluated. 5µm and 10µm sections produced datasets with >500 nuclei for all samples, for 3µm one sample failed. 5µm sections achieved the highest median gene coverage (mRNA expression detected for 835 genes per nucleus), compared with 787 and 792 genes per nucleus for 3 and 10 μm sections, respectively. Given the favorable balance between transcriptome quality and minimal tissue consumption, 5 μm sections were selected as the optimal input material.

We next investigated whether DV200, a commonly used RNA quality metric that may already be available from routine molecular pathology workflows, could predict assay performance (**Figure 2C**). Among the 52 resection specimens, DV200 measurements were available for 37 samples out of the 52. Higher DV200 values were associated with increased nuclei recovery (R = 0.36, p-value = 3.0×10^-2^) and improved gene coverage (R = 0.35, p-value = 3.6×10^-2^), indicating a modest but significant relationship between RNA integrity and sequencing performance. These results suggest that at lower DV200 values, we recover fewer nuclei that exceed the limiting threshold of 500 gene coverage, suspecting a lower quality of the RNA within a fraction of extracted nuclei. Notably, for 82% of samples we could generate a tumor profiling report with over 500 nuclei with over 500 gene coverage across a broad range of DV200 values (0.28-0.95). 90% and 95% success rate were reached for samples above DV200 of 0.60 and 0.81 respectively.

Finally, we evaluated the influence of FFPE block age (**Figure 2D**). FFPE block age was available for 22 specimens (42.3%). For the two most frequent cancer types in this series, bladder and lung, optimal performances - nuclei recovery and coverage - were achieved for the youngest samples. Increasing specimen age was associated with reduced nuclei recovery (R = −0.68, p-value = 4.8×10^-4^) and lower gene coverage (R = −0.49, p-value = 1.9×10^-2^), demonstrating that prolonged storage can negatively affect assay performance. Nevertheless, we could generate successful profiling for 82% of archival specimens ranging from 0 to 11 years old, highlighting the utility of the workflow for retrospective clinical studies. 90% and 95% success-rate were reached for samples younger than 4.9 and 3.2 years old respectively.

In summary, these findings define practical specimen requirements for routine implementation of the assay and demonstrate robust performance across the range of FFPE specimen characteristics encountered in clinical pathology laboratories.

### Cellular composition and cell-type-resolved gene expression are highly reproducible across technical replicates

To evaluate the technical reproducibility of the workflow, we analyzed 20 replicate experiments generated from nine FFPE specimens representing three cancer types. Cellular composition was highly reproducible across technical replicates, with correlations in cell-type proportions ranging from R = 0.88 to 1.00 (median R = 0.994; **Figure 3A–C**). We next evaluated the reproducibility of cell-type-resolved gene expression within malignant cells, the most abundant and clinically relevant population. Average gene expression profiles showed high concordance between replicates (R = 0.745–0.998; median R = 0.982), while the proportion of cells expressing each gene was similarly reproducible (R = 0.746–0.996; median R = 0.958; **Figure 3A–C**).

**Figure 3:**
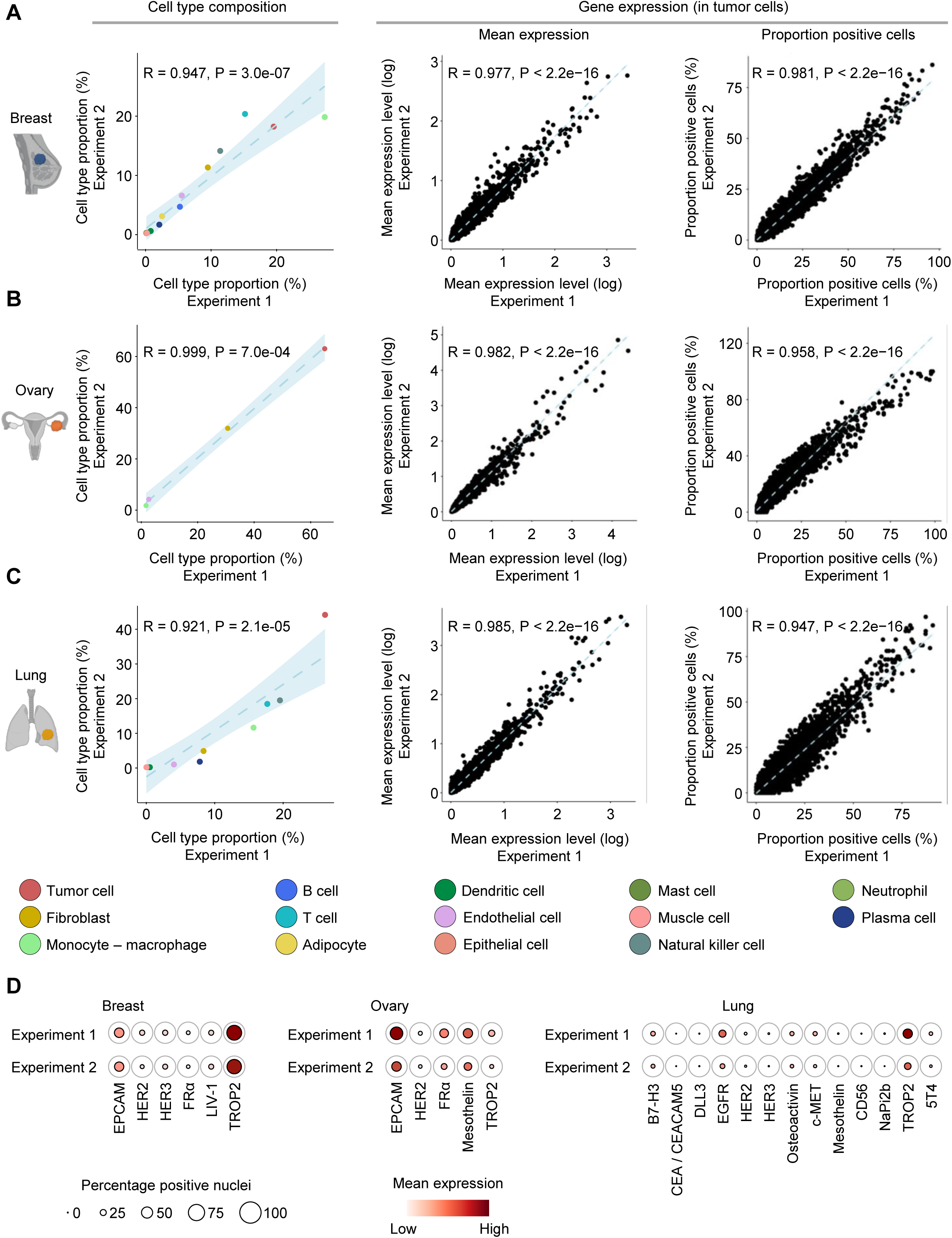
Reproducibility of cellular composition and cell-type-resolved gene expression. (A-B-C) Scatterplot comparing cell composition, and gene expression in tumor cells across two technical replicates for breast, ovarian and lung cancer specimens. Gene expression in tumor cells is assessed at the mean expression level and percent-positive cells. Pearson’s R and corresponding p-values are shown in top left corner. (D) Dot plots displaying expressing levels and percentage of nuclei expressing gene of interest across two technical replicates of the same specimen.

Because quantification of the mRNA expression of therapeutic genomic targets represents a key clinical application of the assay, we next evaluated the reproducibility of clinically actionable biomarkers across tumor types (**Figure 3D**). Consistent cell-type-specific expression patterns were observed for established and emerging antibody-drug conjugate (ADC) targets, including the pan-cancer targets TACSTD2 (TROP2) and EPCAM, the ovarian cancer target FOLR1 (FRα), and the lung cancer targets EGFR and CD276 (B7-H3). Similar reproducibility was observed in fibroblasts, the second most abundant cell population, where average gene expression profiles showed a median correlation of 0.960, while the proportion of positive cells per gene showed a median correlation of 0.935 (**Figure S2**).

Together, these results demonstrate that the workflow reproducibly quantifies both cellular composition and cell-type-resolved gene expression across independent experiments, supporting robust measurement of clinically relevant cellular biomarkers from routine FFPE specimens.

### Routine core biopsies enable cellular profiling of Non-Small Cell Lung Cancer (NSCLC)

We next evaluated whether the workflow could be applied to routine diagnostic core biopsies, which represent one of the most challenging specimen types encountered in clinical practice because of their limited tissue availability. We collected 4 NSCLC lung biopsies as well as 5 biopsies of NSCLC metastatic sites (lymph node, abdomen and pleural cavity).

Despite a median tissue surface area six-fold smaller than surgical resections (5.4 versus 37.0 mm²; **Figure 4A; Figure S3**), routine biopsies generated a total of 7,117 nuclei with a median capture of 585 nuclei per sample and a median coverage of 1,071 genes/nuclei. To increase nuclei recovery for these ultra-low input samples, we lowered the gene-coverage threshold from 500 to 300 genes per nucleus. This increased the total number of retained nuclei from 7,117 to 10,758 (+3,641 nuclei) across nine specimens, corresponding to a median of 1,011 nuclei per sample with a median coverage of 699 genes/nucleus (**Figure 4B**). This change in coverage did not affect our ability to analyze and annotate nuclei (**Figure S4**). OneMap successfully classified 93.7% of these cells, identifying the full spectrum of major NSCLC cellular populations– from the most abundant fibroblast population (10.2%) down to plasma, mast and natural killer cells, which represent 1%, 0.8% and 0.1% of the TME of these biopsies, respectively (**Figure 4C**). Cell-type-resolved gene expression further enabled quantification of clinically relevant therapeutic targets and immune biomarkers, with tumor cells expressing typical ADC target genes such as TROP2 (TACSTD2) and HER2 (ERBB2), and T cells expressing immune checkpoints such as CTLA4 and TIGIT (**Figure 4D**).

**Figure 4:**
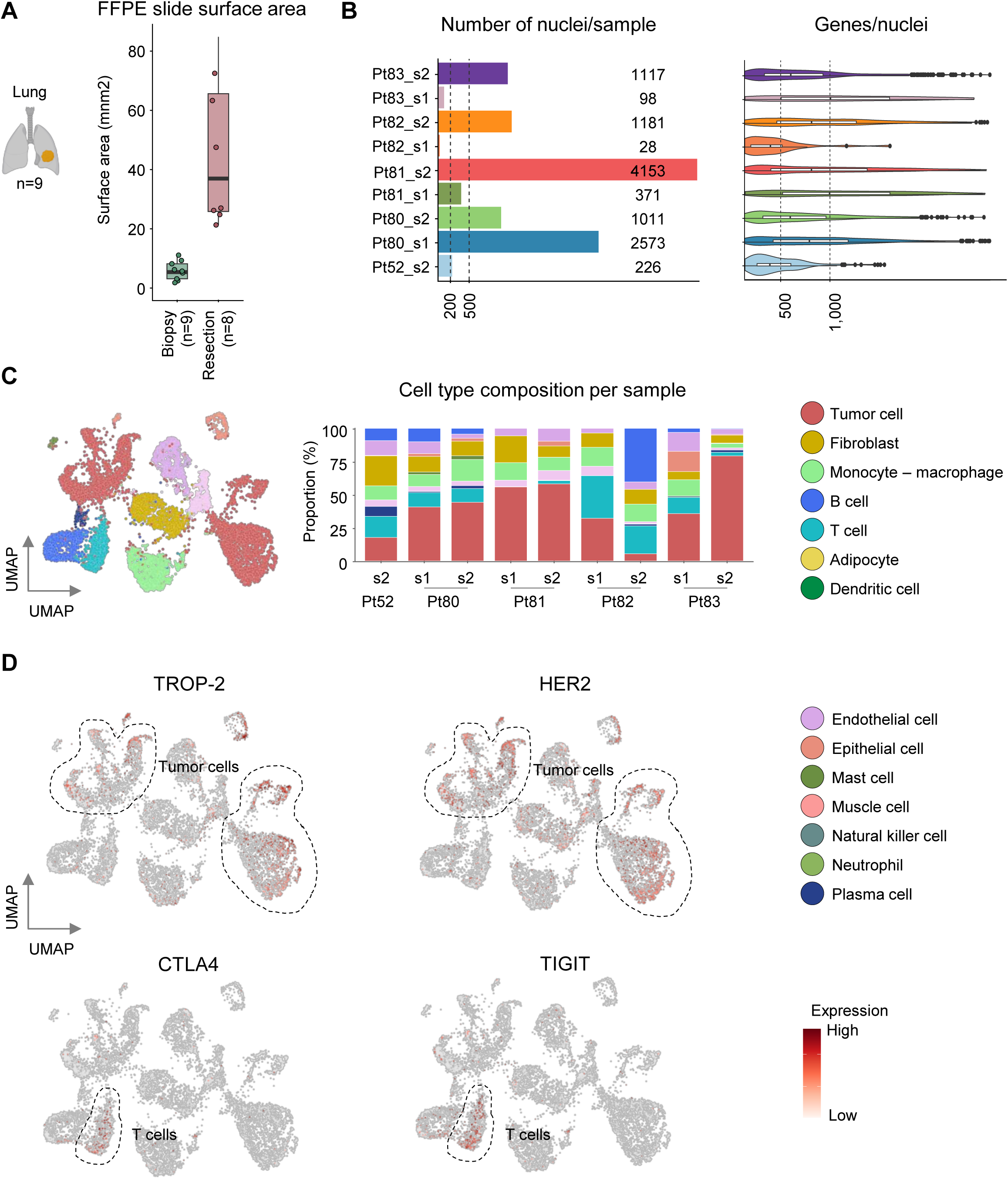
Routine biopsies support clinically informative cellular profiling. (A) Boxplots representing tissue surface area (mm2) of 9 routine diagnostic core NSCLC biopsies and 8 resection specimens. B) Left: bar plots representing numbers of nuclei obtained for each biopsy. Right: violin plots displaying numbers of genes per nuclei for each biopsy. (C) Left: UMAP embedding of the single-nucleus cohort, nuclei are colored according to cell type. Right: bar plot representing cell type composition by patient and sample. (D) UMAP embedding showing gene expression of clinically relevant therapeutic targets and immune biomarkers in tumor cells (top) and T cells (bottom), respectively.

Together, these findings demonstrate that routine FFPE core biopsies are sufficient to generate single-cell transcriptomic profiles, extending the applicability of the workflow to the specimen type most frequently available for prospective molecular characterization of patients with advanced cancer.

### Automated cellular profiling reproduces pathological assessment of tumor immune infiltration

For cellular composition profiling to become clinically informative, measurements generated by the workflow should be highly concordant with pathological assessments routinely used in patient care. We therefore evaluated whether automated quantification of lymphocyte abundance agreed with pathologist-assessed microscope-based tumor-infiltrating lymphocyte counts (TILs) in 16 FFPE triple-negative breast cancer biopsies (**Figure 5**). We gathered samples from 16 patients treated at Institut Curie and generated 702 median nuclei per patient, with 825 detected genes per nucleus (**Figure 5A-B**).

**Figure 5:**
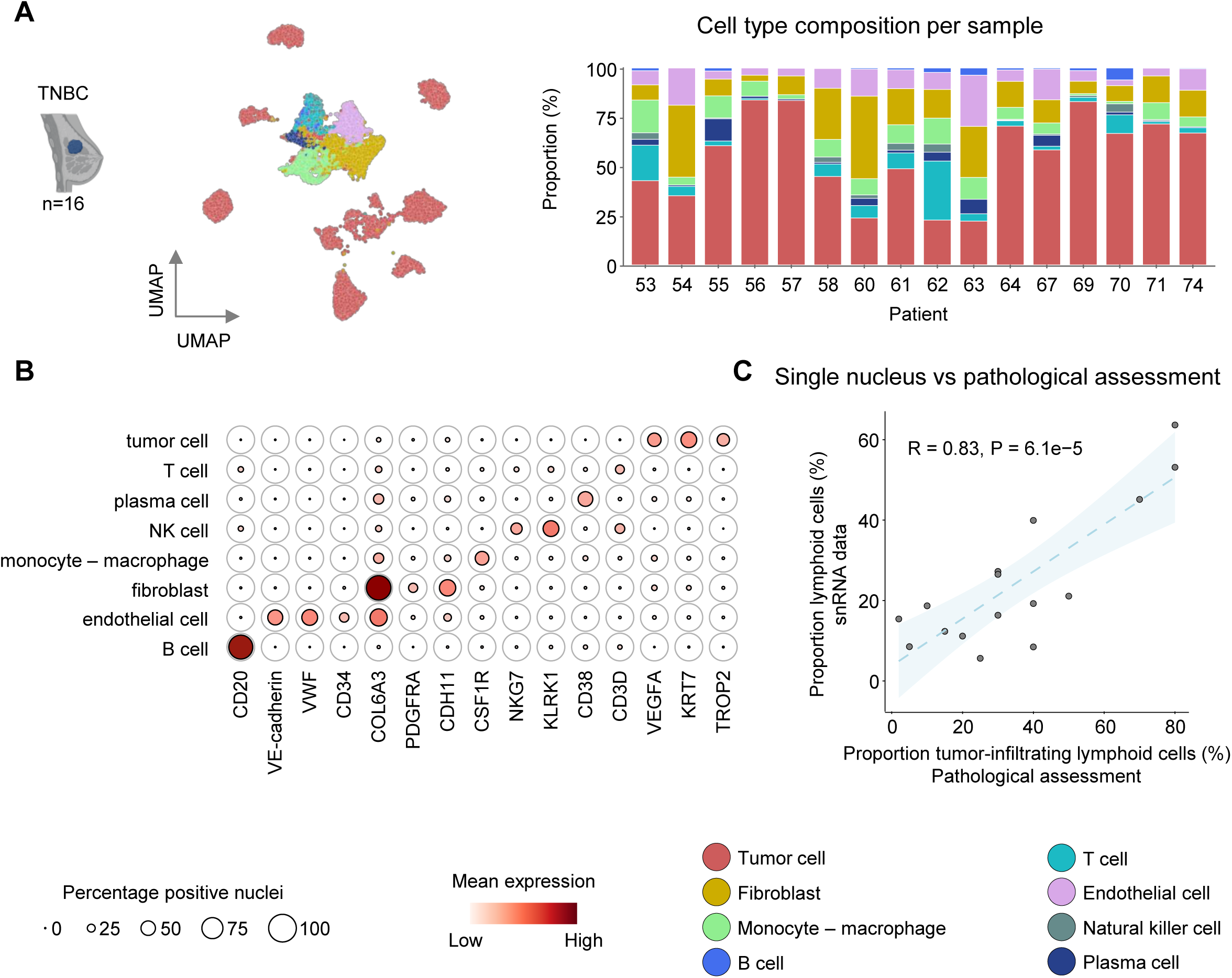
Automated cellular profiling reproduces pathological assessment of immune infiltration. (A) Left: UMAP embedding of the single-nucleus dataset of 16 biopsies in the triple- negative breast cancer (TNBC) cohort, nuclei are colored according to cell type. Right: Barplot representing cell type composition by patient. (B) Dotplots displaying expressing levels and percentage of nuclei expressing cell type marker genes of interest across the cellular populations identified in the cohort. (C) Correlation between transcriptome-derived lymphocyte abundance and pathological TIL assessment. Pearson’s R and corresponding p-values are shown in top left corner.

We observed a strong correlation between pathologist-derived TIL scores and lymphocyte abundance estimated from our automated snRNA-seq data (**Figure 5C**, R = 0.83, p-value = 6.1×10^-5^). Linear regression analysis yielded an intercept of 3.6% and a slope of 0.59, demonstrating a strong quantitative relationship between molecular and histological measurements of immune infiltration. These findings demonstrate that FFPE snRNA-seq faithfully captures biologically and clinically relevant features routinely assessed during pathological evaluation, providing orthogonal validation of the workflow.

### Examples of tumor ecosystem profiling and therapeutic target expression

To illustrate the clinical utility of single-cell profiling from FFPE samples, we examined cell composition and the expression of therapeutic target genes across cell types in four tumors representing distinct cancer types. The first tumor (**Figure 6A**) was a bladder carcinoma (Pt17), comprising 62% tumor cells and a microenvironment dominated by stromal and myeloid cells, with very few T cells (2%) and low expression of immune checkpoint molecules. In contrast, tumor cells highly expressed several ADC targets approved or under clinical investigation in bladder cancer, including NECTIN-4, HER2, and TROP2, the latter showing the strongest and most widespread expression. The second tumor (**Figure 6B**) was an *EGFR*-mutated lung adenocarcinoma (Pt40). Tumor cells co-expressed EGFR and MET, supporting the potential relevance of amivantamab, the clinically approved EGFR/MET bispecific antibody, as a potential therapeutic option. Tumor cells also showed high TROP2 expression, compared to other patients (**Figure S5)**, consistent with eligibility for anti-TROP2 ADC in later treatment lines. The third tumor (**Figure 6C**) was a ER+ breast carcinoma with relatively abundant T- and NK-cell infiltrates. Although PD-1 and PD-L1 expression were low, T cells expressed alternative immune checkpoint molecules, including TIGIT and CTLA4, both under clinical investigation in breast cancer. Tumor cells also expressed targets of approved (TROP2, HER2) and investigational (LIV-1/SLC39A6) ADC, among the highest expression in the cohort (**Figure S5)**. Finally, we analyzed a high-grade serous ovarian carcinoma (HGSOC, Pt32; **Figure 6D**). Limited T-cell infiltration and low PDCD1/CD274 expression were consistent with the generally modest activity of immune checkpoint blockade in this disease. Expression of FOLR1 (FRα), the only approved ADC target in HGSOC, was moderate, whereas the investigational ADC target MSLN was highly expressed by tumor cells. Together, these examples illustrate how single-nucleus RNA-seq from FFPE samples can provide a comprehensive view of the tumor ecosystem, simultaneously characterizing cellular composition, immune contexture, and the cell type-specific expression of therapeutic targets. Across the NSCLC and TNBC cohorts, single-cell profiling revealed marked inter-patient heterogeneity in both the proportion of target-positive tumor cells and target expression levels (**Figure S5**), underscoring the value of single-cell target profiling to refine patient selection for targeted therapies. Such information may complement conventional molecular profiling by supporting the identification of approved targeted therapies and ADCs, as well as potential eligibility for investigational agents in clinical trials.

**Figure 6:**
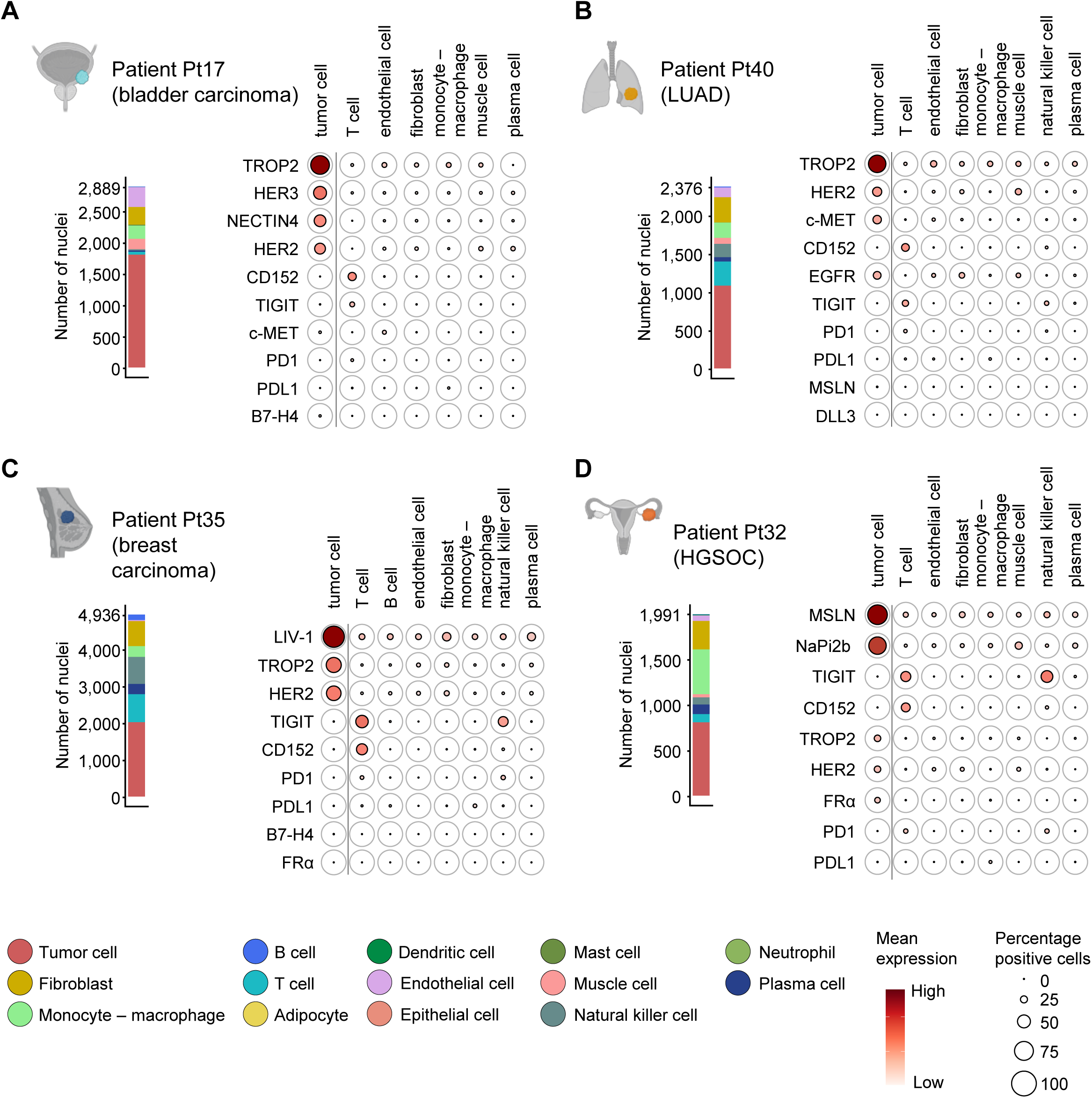
Representative sample-to-report outputs across four cancer types. For each patient, the composition of the tumor ecosystem is shown as a barplot, together with the cell type-specific expression of approved and investigational therapeutic targets, including immune checkpoint molecules and ADC targets, relevant to each cancer type.

## Discussion

In this study, we developed and validated a clinically compatible single-nucleus transcriptomic workflow for routine FFPE specimens. By combining low-input single-nucleus RNA sequencing from four standard 5-μm pathology sections with foundation model-based automated cell annotation and reporting, we established a workflow that operates on individual clinical specimens rather than research-oriented cohorts. Across six major cancer types, the approach demonstrated robust recovery of tumor composition, high technical reproducibility, and concordance with conventional pathological assessment. Importantly, successful profiling was achieved not only in surgical resections but also in smaller core biopsies including archival FFPE material, addressing several practical barriers that have restricted so far the application of single-cell technologies to translational research settings.

A central question for the field is which cellular profiling modalities will ultimately enter clinical oncology workflows. Single-cell platforms are beginning to emerge in clinically oriented laboratories, but accessibility remains limited by specimen requirements, cost, turnaround time, and analytical complexity. The presented FFPE-compatible single-nucleus workflows provide one route to lower these barriers by using the material already available in pathology departments. Other single-cell methods such as spatial transcriptomics, multiplex imaging^24^, or targeted single-cell sequencing^25^ and liquid biopsy approaches may also each address complementary clinical questions. Clinical implementation is for example emerging for targeted single-cell DNA sequencing, which is being incorporated into prospective studies to resolve clonal architecture and therapeutic resistance in hematological malignancies, complementing transcriptomic approaches that quantify cellular composition and functional cell states. Rather than competing modalities, these approaches may converge toward shared data models in which clinically relevant cell states, immune and stromal populations, or multicellular “ecotypes” are defined in tissue and then monitored through complementary assays, including liquid biopsies^26^.

The next step is to establish clinical utility. Several studies now suggest that cellular composition and cell-state information at baseline can inform diagnosis, treatment response, or resistance^15^. Baseline tumor cellular composition is emerging as a predictor of treatment response trajectories across multiple cancer types, including bladder^27^, TNBC^28^, lung cancer^29^. Similar concepts are emerging in ovarian cancer, where transcriptomic and epigenomic profiling suggests that tumors resistant to standard of care platinum-based chemotherapy can be identified and “primed” before therapy^6^. Together, these studies support the idea that single-cell transcriptome profiling can identify clinically relevant features at baseline beyond conventional genomic or pathology assays, but prospective studies will be required to show whether these measurements improve patient stratification, treatment selection, or disease monitoring.

A major challenge for the field is the transition from academic single-cell technologies to regulated clinical assays. As previously observed for next-generation sequencing ^30^, widespread adoption will require standardized workflows^31^, quality control metrics, inter-laboratory benchmarking, and accreditation within existing frameworks such as CLIA (Clinical Laboratory Improvement Amendments), CLEP (Clinical Laboratory Evaluation Program; New York State Department of Health Wadsworth Center) and CAP (College of American Pathologists). Benchmarking single-cell technologies presents unique challenges because the output extends beyond molecular measurements to include cell identities, cell states, and TME-level features. Future efforts will therefore need to establish reproducible standards for cell-type assignment, target quantification, and ecosystem characterization, using orthogonal approaches such as pathology, flow cytometry, and multiplex imaging as reference frameworks. Ultimately, clinical adoption will depend on demonstrating that single-cell profiling improves patient stratification, treatment selection, or disease monitoring. The development of clinically compatible sample-to- report workflows operating on routine FFPE specimens represents an important step toward this goal.

---

**Supplementary Figure S1.**
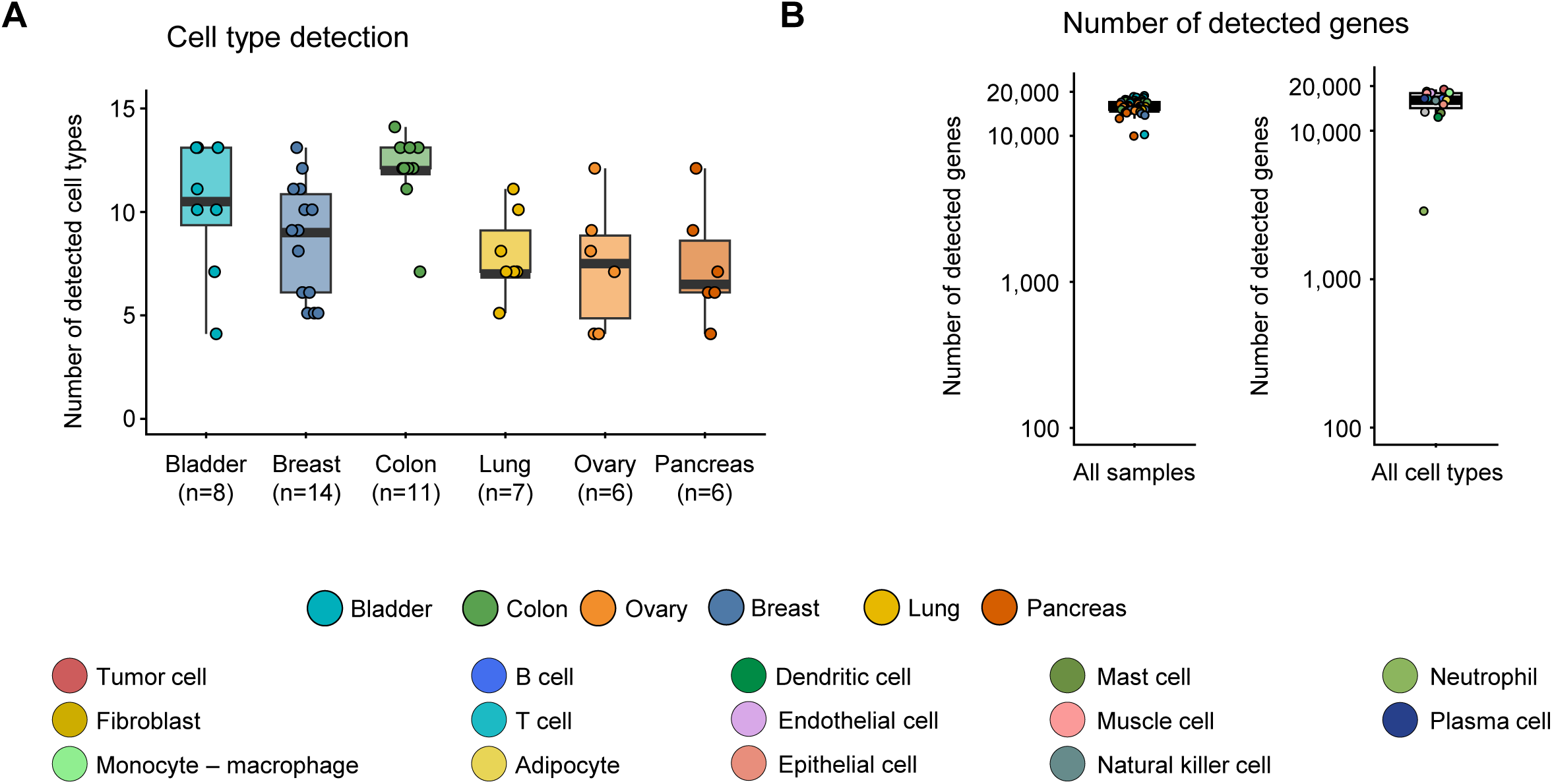
(A) Boxplots representing numbers of cell types detected in each resection sample (n=52), grouped by cancer types. (B) Left: boxplot of number of detected genes across all 52 resections, samples are colored according to cancer type. Right: boxplot of number of detected genes across all 14 detected cell types, points are colored by cell type.

**Supplementary Figure S2.**
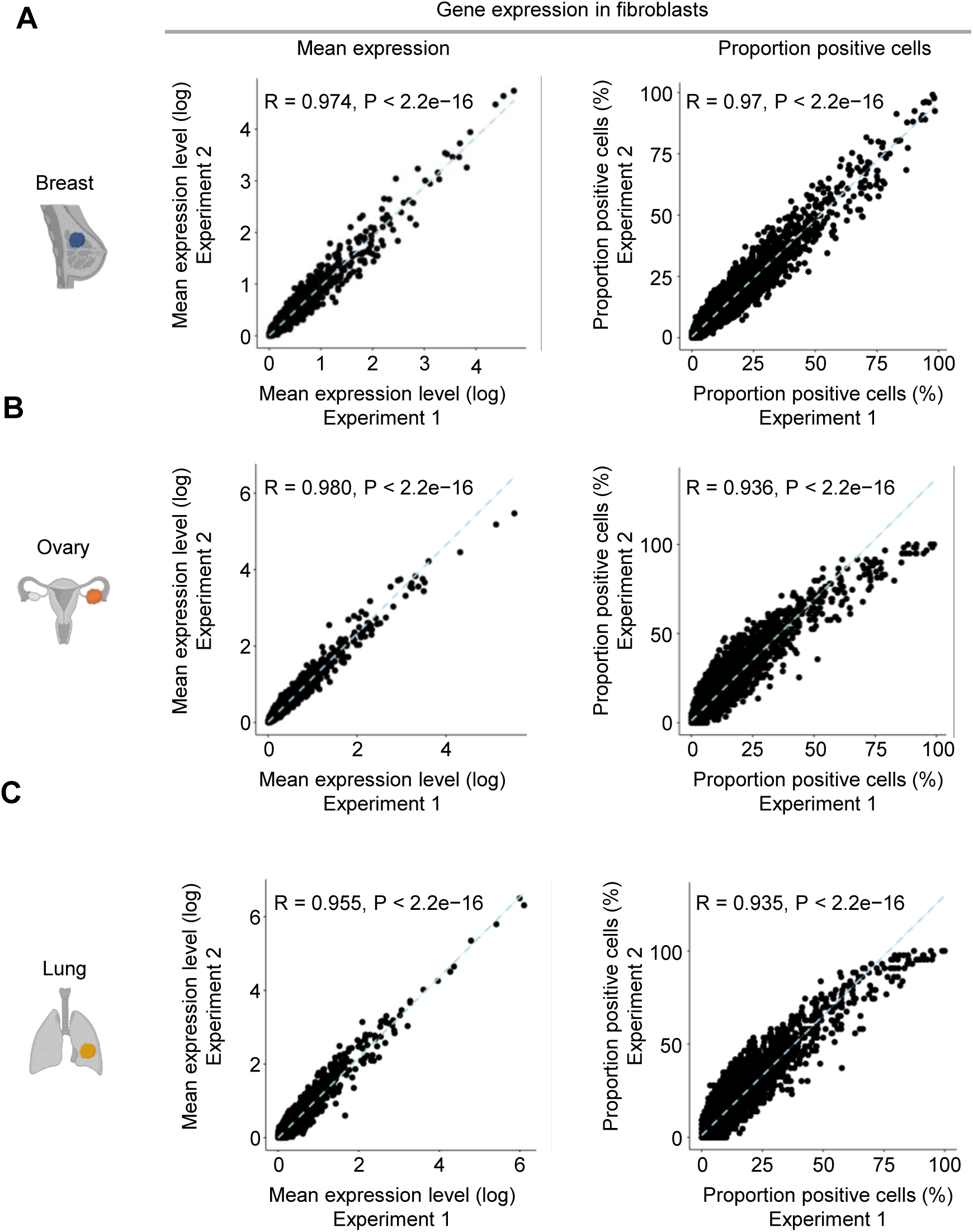
(A-B-C) Scatterplot comparing gene expression level and percent-positive cells in fibroblasts across two technical replicates for breast, ovarian and lung cancer specimens. Pearson’s R and corresponding p-values are shown in top left corner.

**Supplementary Figure S3.**
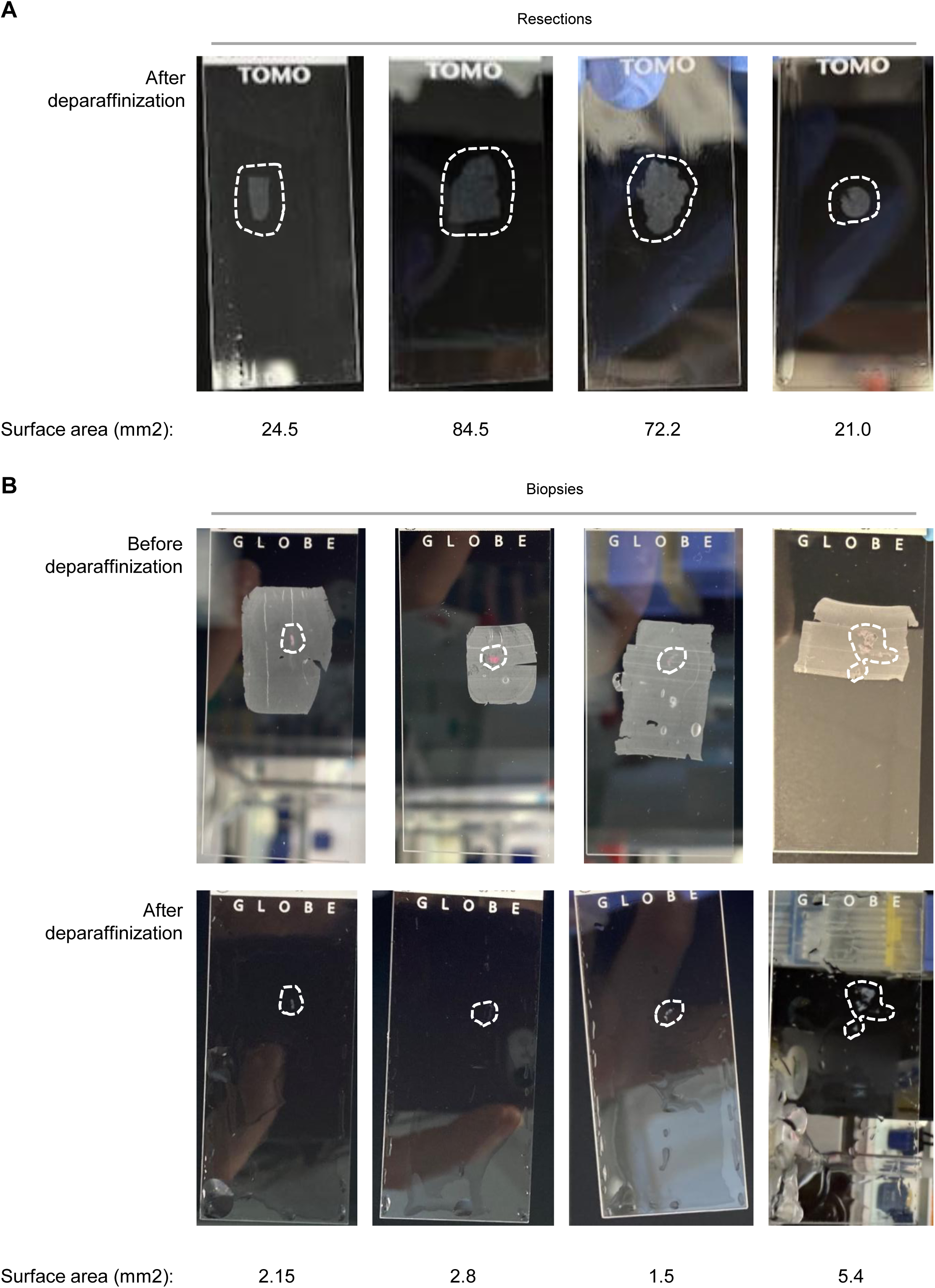
(A-B) Four representative FFPE slides and their corresponding surface areas from resections and biopsy specimens, as shown in Figure 4A. For biopsy specimens, views before and after paraffinization are provided to more clearly visualize specimen positioning.

**Supplementary Figure S4.**
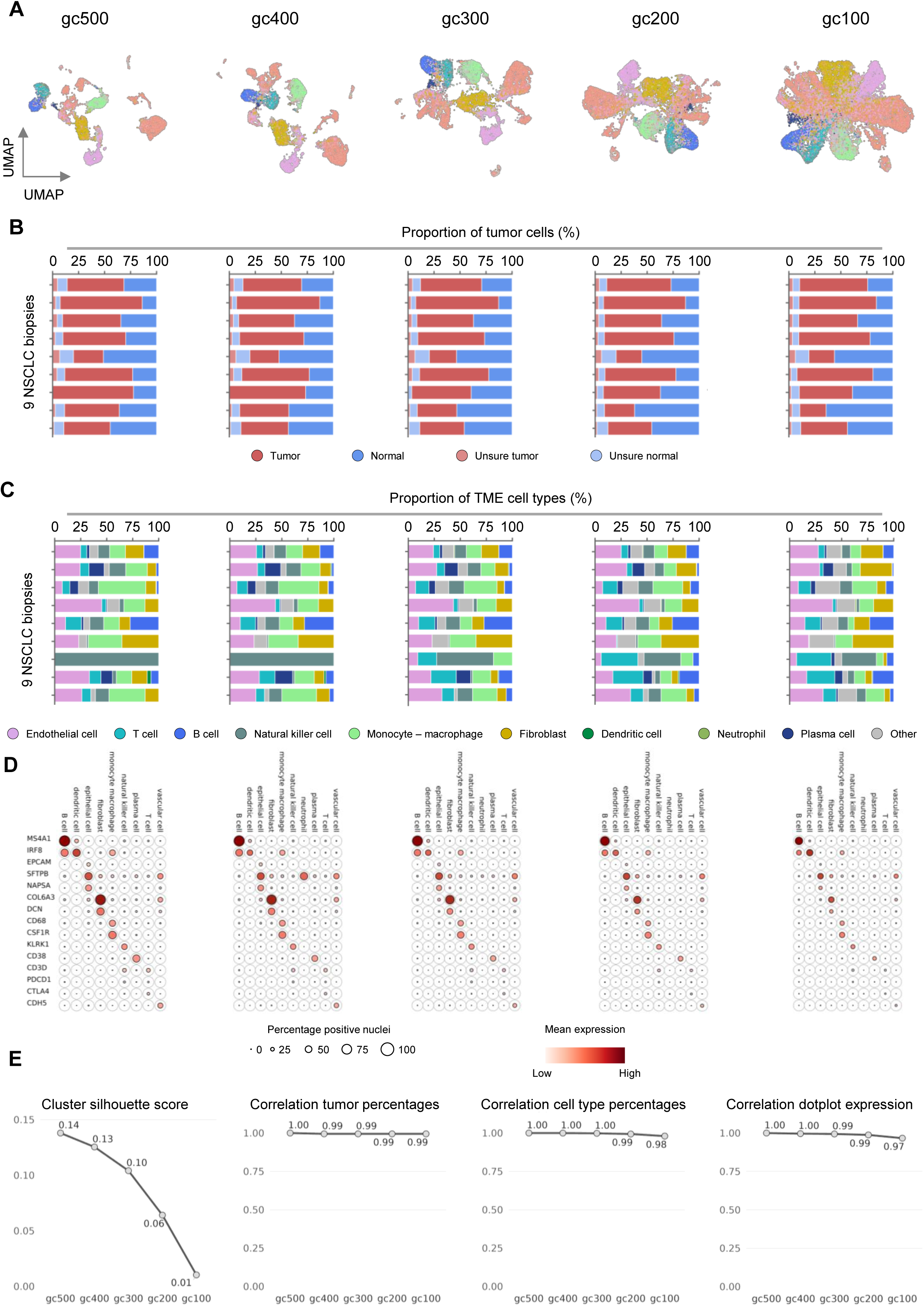
(A) UMAP embedding of the single-nucleus dataset of 9 NSCLC biopsies using 5 different thresholds for gene coverage (gc500, genes/nucleus > 500; gc400, genes/nucleus > 400; gc300, genes/nucleus > 300; gc200, genes/nucleus > 200; gc100, genes/nucleus > 100), nuclei are colored according to cell type. (B) Bar plots representing tumor cell composition by sample. (C) Bar plots representing TME cell type composition by sample. (D) Dotplots displaying expressing levels and percentage of nuclei expressing cell type marker genes of interest across the cellular populations identified in the cohort. (E) Line plots summarizing the evolution of the quality metrics when varying the threshold for gene coverage. Metrics include the cluster silhouette score for clustering resolution 0.8, the correlation of tumor percentages, the correlation of cell type percentages and the correlation of dotplot expression.

**Supplementary Figure S5.**
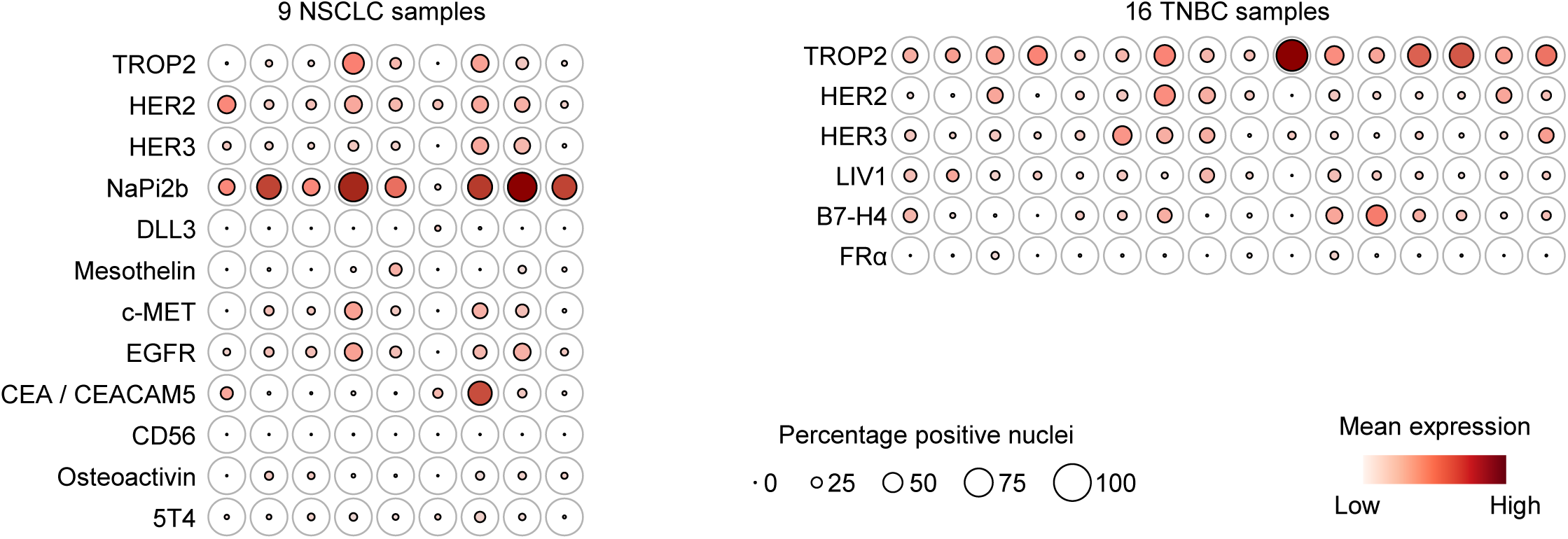
(A-B) Dot plots displaying expressing levels and percentage of nuclei expressing of approved and investigational therapeutic targets of ADCs and multispecific antibodies, relevant to the cancer type, across tumor cells of each of the 9 NSCLC biopsies (A) and of the 16 TNBC biopsies (B).

## Material and methods

### Patient samples

Formalin-fixed paraffin-embedded (FFPE) tumor specimens were obtained from patients treated at Institut Curie (Paris, France), Centre Léon Bérard (Lyon, France), Hôpital Bichat–Claude Bernard (Paris, France), and Memorial Sloan Kettering Cancer Center (New York, USA). Samples included both surgical resection specimens and diagnostic core biopsies collected as part of routine clinical care. The use of human specimens for this study was approved by the appropriate institutional review boards or ethics committees at each participating institution (Institutional Review Board No. DATA250199 and No. 00006477). Written informed consent for research use of tissue was obtained from all patients when required by local regulations. Where applicable, the use of archived specimens was approved under institutional protocols, permitting research on residual clinical material. The study was conducted in accordance with the principles of the Declaration of Helsinki and all applicable national and institutional regulations.

### Nuclei preparation from FFPE tumor sections

For single-nucleus RNA sequencing (snRNA-seq), formalin-fixed paraffin-embedded (FFPE) surgical specimens or biopsies were sectioned at 5 µm thickness, and four slides were processed per patient. Nuclei were isolated as in Landais et al^6^. Immediately after nuclei isolation, snRNA-seq libraries were generated using the Chromium GEM-X Flex Gene Expression v2 platform (10x Genomics) according to the manufacturer’s instructions. Following GEM generation, cDNA amplification and library construction were performed according to the manufacturer’s protocol. Library quality and concentration were assessed using standard quality control procedures (Qubit and Tapestation) prior to paired-end sequencing on an Illumina NextSeq 2000 platform.

### From raw data to count matrices

We used a custom workflow to process raw FASTQ files. The UMI count matrices were generated using *Cell Ranger* with the human genome reference (GrCh38/hg38). We used DoubletCollection to detect doublet cells with *doubletCells*, *cxds*, *bcds*, *hybrid*, *scDblFinder*, *Scrublet*, *DoubletDetection* and *DoubletFinder* algorithms. Each cell detected as doublet by ≥3 methods was removed. We retained only high-quality cells (>500 genes detected and <20% of mitochondrial RNA reads; for the NSCLC biopsies, high-quality cells were selected using >300 genes detected and <20% of mitochondrial RNA reads). Counts for each cell were divided by the total counts for that cell, multiplied by a scale factor of 10,000, and subsequently log1p- normalized using *Seurat*’s *NormalizeData* function. We then applied *Seurat*’s default workflow, including dimensionality reduction with PCA and UMAP, followed by clustering using the Leiden algorithm.

### Target gene quantification

Dot plots display the genes’ mean expressing levels and percentages of gene-positive nuclei across two or more cell groups (technical replicates, samples or cell types). Mean expression was calculated as the mean normalized expression across the cell group. The proportion of gene-positive cells is defined as the fraction of cells with at least one UMI assigned to the gene’s transcript.

## Data availability

Processed single-nucleus RNA sequencing data supporting the findings of this study are available from the corresponding authors upon request, subject to applicable ethical and institutional approvals. All other data supporting the findings of this study are available within the Article and its Supplementary Information.

## Author contribution

J.W., J.B. and M.G. contributed equally to this work. E.L. and C.V. jointly conceived and supervised the study. J.B. and M.G. developed the experimental workflow and performed the experiments. J.W., B.S., S.G., A.R., M.D.-R. and E.A. developed the OneMap computational workflow and performed the computational analyses. A.P., H.Y., J.-Y.B., C.G., L.N.-B., A.V.S., L.V., E.X., L.C. and J.S.R. provided clinical samples and clinical and pathological expertise. V.M. contributed to study design and interpretation of the results. J.W., E.L. and C.V. wrote the manuscript with input from all authors. All authors reviewed, revised and approved the final manuscript.

## Acknowledgements.

We thank all members of the One Biosciences team for scientific discussions. We thank Christine Sheenan for critical reading of the manuscript.

## Competing interests

CV is a co-founder of One Biosciences. The authors declare that patents have been filed related to the work reported in this paper.

## Notes

### Summary of Updates

A mistake in one author name

